# Role of Glycogen Metabolism in *Clostridioides difficile* Virulence

**DOI:** 10.1101/2024.04.15.589614

**Authors:** Md Kamrul Hasan, Marjorie Pizzarro-Guajardo, Javier Sanchez, Revathi Govind

## Abstract

Glycogen plays a vital role as an energy reserve in various bacterial and fungal species. *Clostridioides difficile* possesses a glycogen metabolism operon that contains genes for both glycogen synthesis and utilization. In our investigation, we focused on understanding the significance of glycogen metabolism in the physiology and pathogenesis of *C. difficile*. To explore this, we engineered a *C. difficile* JIR8094 strain lacking glycogen synthesis capability by introducing a group II intron into the *glgC* gene, the operon’s first component. Quantification of intracellular glycogen levels validated the impact of this modification. Interestingly, the mutant strain exhibited a 1.5-fold increase in toxin production compared to the parental strain, without significant changes in sporulation rate. Our analysis also revealed that wild-type *C. difficile* spores contained glycogen, whereas spores from the mutant strain lacking stored glycogen showed increased sensitivity to physical and chemical treatments and had a shorter storage life. By suppressing *glgP* expression, the gene coding for glycogen-phosphorylase, via CRISPRi, we demonstrated that glycogen accumulation but not the utilization is needed for spore resilience in *C. difficile*. Transmission Electron Microscopy (TEM) analysis revealed a significantly lower core/cortex ratio in *glgC* mutant strain spores. In hamster challenge experiments, both the parental and *glgC* mutant strains colonized hosts similarly; however, the mutant strain failed to induce infection relapse after antibiotic treatment cessation. These findings highlight the importance of glycogen metabolism in *C. difficile* spore resilience and suggest its role in disease relapse.

**Importance:** This study on the role of glycogen metabolism in *C. difficile* highlights its critical involvement in the pathogen’s energy management, its pathogenicity and resilience. Our results also revealed that glycogen presence in spores is pivotal for their structural integrity and resistance to adverse conditions, which is essential for their longevity and infectivity. Importantly, the inability of the mutant strain to cause infection relapse in hamsters post-antibiotic treatment pinpoints a potential target for therapeutic interventions, highlighting the importance of glycogen in disease dynamics. This research thus significantly advances our understanding of *C. difficile* physiology and pathogenesis, offering new avenues for combating its persistence and recurrence.

## Introduction

In prokaryotic cells, Glycogen works as a storage carbohydrate energy source and provides sugar molecules during nutrient deprivation [1]. Glycogen is an excellent storage carbohydrate since it makes dense granules with minimum effect on the osmolarity of the cell. The synthesis of glycogen is highest in bacteria when excess carbohydrate is available [2]. Glycogen metabolism has been linked to virulence in several pathogens. Examples include sporulation in *Bacillus subtilis* [3], biofilm formation, and virulence in *Salmonella enteritidis* [4]. In *Vibrio cholera*, stored glycogen is responsible for increased environmental persistence and transmission [5]. In addition to the virulence properties, glycogen biosynthesis has significant effects on the physiology of the bacteria too. In *Lactobacillus acidophilus*, glycogen metabolism impacts gut retention and stress tolerance [6]. In an *in vivo* mouse model study, *E. coli* strains with disrupted glycogen biosynthesis genes showed a substantial reduction in gut colonization capacity [7], indicating the significance of glycogen metabolism for commensal bacteria survivability.

Studies have shown that *C. difficile* glycogen operon is directly under the transcriptional control of the virulence master regulator CodY [8]. Given the significance of CodY regulation in *C. difficile* virulence per nutrient availability, we hypothesized that glycogen metabolism is tightly linked with the *C. difficile’s* nutrient signaling and disruption of glycogen metabolism could have a significant effect on the virulence properties in *C. difficile*. Additionally, a study on the proteomics of a *C. difficile Δspo0A* mutant showed proteins of the Glycogen metabolism pathway to be downregulated in this mutant, hinting at a possible connection between sporulation and glycogen metabolism [9].

Three primary enzymes contribute to the biochemical pathway of glycogen biosynthesis in bacteria. First, the substrate for glycogen, which in the case of bacteria is Glucose -1-Phosphate, gets converted to ADP-Glucose (ADPG) by the enzyme ADP-Glucose Pyrophosphorylase encoded by *glgC* [2]. Using ADPG as a sugar donor, glycogen is synthesized by glycogen synthase-encoded by *glgA*. After chain elongation by *glgA*, glycogen branching enzyme (*glgB*) catalyzes the formation of branched oligosaccharide chains having α-1,6-glucosidic linkages. For the glycogen utilization, the enzyme glycogen phosphorylase coded by *glg*P is responsible for breaking down glycogen to glucose-1-phosphate. In *C. difficile,* the glycogen operon is constituted of 5 genes: *glgC, glgD, glgA, glgP,* and putative alpha-amylase (Figure 1, Supplemental Figure 1). The *glgB*, coding for the glycogen branching enzyme is not part of the operon, but located in a different location in the genome. *B. subtilis* has a similar assortment of genes in its glycogen operon, albeit in a different organization. Many other gut-living and pathogenic bacteria have a comparable assortment of *glg* genes in various orders (Supplemental Figure 1).

**Figure 1.**
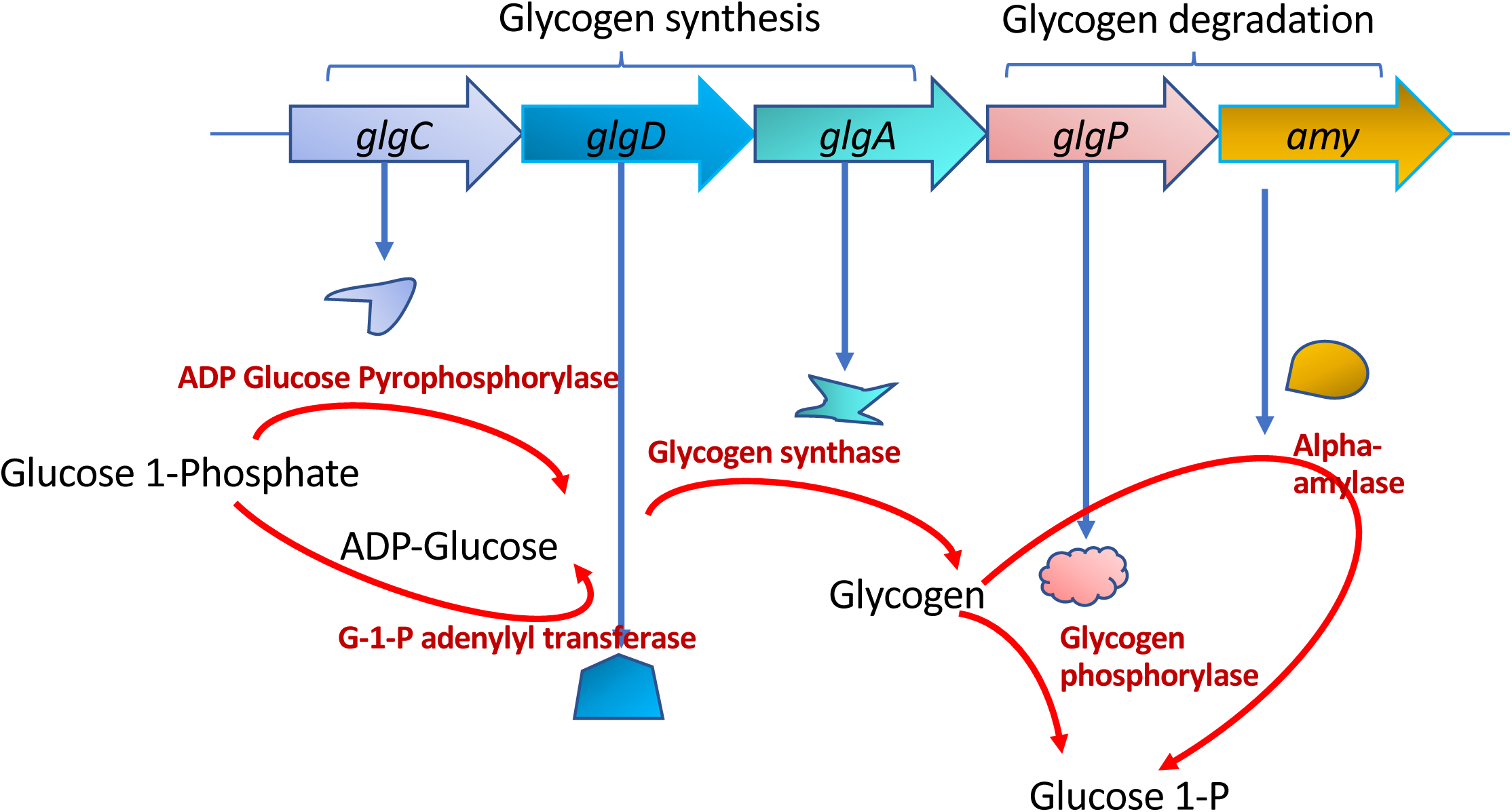
*C. difficile* Glycogen operon schematics. Enzymes in glycogen metabolism pathway and the respective genes (*glgC, glgD, glgA, glgP*, and *amy*) encoding them. The substrates and products of the enzymes are also illustrated with arrows denoting the direction of enzymatic reactions. *glgC, glgD,* and *glgA* encodes enzymes of glycogen synthesis pathway, whereas *glgP* and *amy* encodes enzymes of glycogen degradation pathway. Gene *glgB* coding for the essential glycogen-branching enzyme is not part of the glycogen operon and thus not shown here.

In this investigation, our aim was to elucidate the influence of glycogen metabolism on various virulence traits of *C. difficile*. To achieve this, we engineered a mutant *C. difficile* strain (JIR8094:: *glgC*) with impaired glycogen metabolism and conducted a comparative analysis to assess the importance of glycogen in *C. difficile* physiology and virulence. We present and discuss the results of our findings below.

## Materials and Methods

### Bacteria strains and growth conditions

*C. difficile* WT (JIR8094) and the mutant strain (JIR8094::*glgC*) were grown anaerobically in TY (Tryptose, Yeast Extract), liquid, or TY with 0.5% glucose (TYG) or 70:30 medium as described previously [10,11]. Kanamycin (50 µg/ml), Thiamphenicol (15 µg/ml), and Lincomycin (Lin; 20 µg/ml) were supplemented whenever necessary. *Escherichia coli* strains were grown in (LB) broth. *E. coli* strain S17 [12], carrying the Clostron plasmic construct used for conjugation was grown in LB supplemented with chloramphenicol (25 µg/ml) as required.

### Construction of a *C. difficile* JIR8094:: *glgC* mutant

The *C. difficile* JIR8094:: *glgC* mutant strain was constructed through the utilization of the ClosTron mutagenesis system [13]. The specific insertion site for the group II intron, oriented in the antisense direction of the *glgC* ORF, was determined using the Perutka algorithm, an online design tool accessible at http://www.clostron.com. Subsequently, the intron with the retargeted sequence was synthesized through the Genewiz gene synthesis service and cloned into the pMTL007-CE5 vector. This resultant plasmid, designated as pMTL007-CE5::Cdi-*glgC*, was introduced into the JIR8094 strain through conjugation, as previously described [10,14,15]. Transconjugants exhibiting resistance to thiamphenicol were then plated on TY agar supplemented with lincomycin (15 µg·ml−1). This procedure aimed to identify colonies harboring ltrB (Ll.ltrB) intron insertions within the targeted *glgC* gene located on the chromosome of the JIR8094 strain. To confirm the presence of the *glgC* mutant, PCR was conducted using primers specific to *glgC* (Supplemental table 1) in conjunction with the intron specific EBS universal primers (Supplemental Figure 2).

### Growth comparison

Growth comparisons of WT and the mutant were made using both the spectrophotometric and colony counting methods. The cells were grown for a period of 30 hours or more in an anaerobic chamber in TY and TYG medium. To gauge cell density, OD600 measurements were obtained at specified time intervals. Simultaneously, an equivalent volume of cells from both the WT and the mutant strains was subjected to serial dilution using fresh TY media and were plated onto TY agar plates and allowed to grow for 24 hours. The resulting colonies were enumerated and graphed to derive the colony-forming units (CFU) count per milliliter.

### Toxin assay

Cytosolic toxins were measured using toxin ELISA following established methods [10]. Specifically, 1 ml of the 16-hour cultures of both WT and the JIR8094::*glgC* mutant strains were collected. These samples were then suspended in 200 μl of sterile PBS, followed by sonication and centrifugation to extract the cytosolic proteins. Subsequently, 100 μg of these cytosolic proteins were employed for assessing toxin levels through the utilization of the *C. difficile* premier Toxin A & B ELISA kit, provided by Meridian Diagnostics Inc. (Cincinnati, OH).

### Sporulation assay (ethanol resistance method)

*C. difficile* strains were inoculated into and grown on 70:30 sporulation agar as described previously [11]. After 30 h of growth, cells were scraped from the plates and suspended in 70:30 sporulation liquid medium to an OD_600_ of 1.0. Cells were immediately serially diluted and plated onto TY agar–0.1% taurocholate to enumerate viable vegetative cells and spores. A 0.5-ml aliquot of the culture was removed from the chamber, mixed with 0.5 ml of 95% ethanol, subjected to vortex mixing, and incubated at room temperature for 15 min. Ethanol-treated cells were serially diluted in PBS, returned to the anaerobic chamber, and plated onto TY agar–0.1% taurocholate plates to enumerate spores. After 24 h of growth, CFU were enumerated, and percent sporulation was calculated as the number of ethanol-resistant spores divided by the total number of viable cells (vegetative cells and spores).

### Spore preparation

*C. difficile* spores were prepared as described earlier [16], but with minor modifications as follows*. C. difficile* strains were streaked onto 20 plates of 70:30 agar medium. After 4 days, the growth from each plate was scraped into 1 ml of sterile water and incubated overnight at 4 °C. The following day, the growth suspension was washed five times with sterile water. In between each wash step, the layer of white cell debris was removed. After washing, the spores/cell debris suspension in water was layered on top of a 60% (w / v) sucrose solution. The gradient was centrifuged in a swinging-bucket rotor at 3,200 X g for 20 minutes. During the centrifugation, the dense spores travel through the sucrose and form a pellet on the bottom while the cell debris is caught in the upper layer. After centrifugation, the solution was removed and the pellet containing the spores was washed five times with sterile water to remove any sucrose. The spores were then suspended in sterile water up to 1 ml. Purified spores were examined under phase contrast microscopy to determine the quality of the preparation.

### Spore sensitivity assay

For the heat resistance assessment, approximately 1 × 10^5^ purified spores, prepared as detailed earlier, were suspended in 500 µl of water and subjected to incubation at temperatures of 70°C and 90°C. For the evaluation of sensitivity to various chemical agents, highly concentrated spores (1 × 10^5^ in 5 µl of water) were resuspended in 495 µl of disinfecting solutions. Disinfectants tested, includes 10% and 1% bleach, 3% hydrogen peroxide (H_2_O_2_), 1N sulfuric acid, and 1N sodium hydroxide (NaOH). Sampling was conducted at intervals of 5, 10, 30, and 60 minutes. Subsequently, the collected samples were serially diluted in phosphate-buffered saline (PBS), plated onto TY agar plates supplemented with 0.1% taurocholate, and cultivated anaerobically for a duration of 48 hours. Untreated spores were serially diluted and were plated onto TY agar containing 0.1% taurocholate before the start of the experiment. The percentage of viable spores was determined by dividing the colony-forming units (CFU) count from the treated samples by the count obtained from the untreated samples, and then multiplying the result by one hundred.

### Quantitative analysis of intracellular glycogen content

The intracellular level of glycogen was measured using the glycogen assay kit (SIGMA MAK 016) using the manufacturer’s instruction. Briefly, *C. difficile* was grown in TY medium and were harvested at different timepoints. Then they were washed with PBS for three times, and the pellet was stored at -80°C until the assay is performed. Cell pellets were resuspended in PBS and were sonicated for 15 seconds, three times and the supernatants were collected. To measure the glycogen in the spores, purified spores were resuspended in PBS and were sonicated for 15 seconds, ten times before collecting the supernatants. We measured the protein content of the lysates using Bradford assay.

For the glycogen measurement, 10 µL of the supernatants were transferred into individual wells of a 96-well black walled clear bottom assay plate and 15 µL of hydrolysis buffer was added to each well, resulting in a final volume of 25 µL. Then, 1 µL of the hydrolysis enzyme mix (amylo glucosidase) was added to the wells, mixed gently before incubating the plate in a dark environment at room temperature for 30 minutes. Immediately following the incubation, 25 µL of the development enzyme mix (hexokinase/glucose-6-phosphate dehydrogenase) was added to the wells. Fluorescence intensity was measured at the excitation wavelength (λex) of 535 nm and the emission wavelength (λem) of 587 nm. A standard curve was generated using the known amount of glycogen. Glycogen measured were presented as µg per mg of total protein content. Glycogen in spores were measured following the same procedure as described above. We enumerated the number of spores in the test sample before glycogen measurement and the glycogen content of spores are presented as ngs/10^8^ spores.

### Transmission Electron Microscopy

For transmission electron microscopy, *C. difficile* spores (10^10^) were fixed overnight with 3% glutaraldehyde, 0.1 M cacodylate buffer (pH 7.2) at 4°C. Samples were centrifuged at 14k rpm for 5 minutes and supernatant discarded. They were then stained with 1% osmium tetroxide in 0.05M HEPES buffer (pH 7.4) overnight at 4°C. Treated samples were washed 5x with distilled water. The spores were dehydrated stepwise in 30%, 50%, 70%, 90% for 15 minutes respectively, followed by dehydration with 100% acetone 3 times for 30 minutes at each step. Spore samples were embedded in modified Spurr’s resin (Quetol ERL 4221 resin; EMS; RT 14300) in a Pelco Biowave processor (Ted Pella, Inc.) as described earlier [17]. Ultrathin sections ∼100nm (silver-gold color) were obtained using Leica UC7 Ultramicrotome and placed on glow-discharged carbon coated 300-mesh Cu grids. Grids were double lead stained with 2% uranyl acetate for 5 minutes in the dark and washed with filter sterilized distilled water followed by 5 minute staining with Reynold’s lead citrate and washed as described. All ultrathin TEM sections were imaged on a JEOL 1200 EX TEM (JEOL, Ltd.) at 100 kV, and images were recorded on an SIA-15C charge-coupled device (CCD). All equipment used is located at the Texas A and M University Microscopy and Imaging Center Core Facility (RRID: SCR_022128).

All measurements were made with ImageJ, in nanometers. The core diameter was measured 3 times. Average was calculated and divided by 2 to obtain the radius (r). Then, all layers (HP: hair projection, Ex: exosporium, OC: outer coat, IC: inner coat, Cx: Cortex, GCW: germinal cell wall) were measured 3 times. In the case of oval cuts of the spores, the 3 core measurements presented high variability. In these cases, the lower measure was kept, indicating the smallest core diameter should represent the diameter if the cut would give a circle. Radius of the cortex was measured by adding the average of thickness of cortex (from IM: “inner membrane to Cx”)+ r (core). Statistical analysis was made with t test and Mann-Whitney test, comparing wild type and the JIR8094::*glgC* mutant.

### *In vivo* animal model study for *C. difficile* pathogenesis

Male and female Syrian golden hamsters (100–120 g) were used for *C. difficile* infection. Upon their arrival, fecal pellets were collected from all hamsters, homogenized in 1 ml saline, and examined for *C. difficile* by plating on CCFA-TA (Cycloserine Cefoxitin Fructose Agar-0.1% Taurocholate) to ensure that the animals did not harbor endogenous *C. difficile*. After this initial screen, they were housed individually in sterile cages with ad libitum access to food and water for the duration of the study.

For the primary infection model, hamsters were first gavaged with 30 mg/kg clindamycin [18]. *C. difficile* infection was initiated three days after clindamycin administration by gavage with *C. difficile* spores. Bacterial spores used as inoculums were standardized and prepared just a few days before the challenge. Immediately before and after infecting the animal, a 10 μL sample of the inoculum was plated onto TY agar with cefoxitin to confirm the spore count and viability. There were three groups of animals, including the uninfected control group. Eight animals per group were used for the infection. Approximately, 2000 spores of *C. difficile* JIR8094 or *C. difficile* JIR8094:: *glgC* were used for the animal challenge. In the uninfected control (group 3), only six animals were used, and they received only antibiotics and sterile PBS. Animals were monitored for signs of disease (lethargy, poor fur coat, sunken eyes, hunched posture, and wet tail) every four hours (six times per day) throughout the study period. Hamsters were scored from 1 to 5 for the signs mentioned above (1-normal and 5-severe). Hamsters showing signs of severe disease (a cumulative score of 12 or above) were euthanized by CO^2^ asphyxiation. Surviving hamsters were euthanized 15 days after *C. difficile* infection. The survival data of the challenged animals were graphed as Kaplan-Meier survival analyses and compared for statistical significance using the log-rank test using GraphPad Prism 7 software (GraphPad Software, San Diego, CA).

To evaluate the recurrence of *C. difficile* disease, a relapse model was employed in hamsters. Initially, the hamsters were administered Clindamycin and subsequently challenged with *C. difficile*, mirroring the primary infection model. However, 24 hours following the *C. difficile* challenge, the hamsters received daily vancomycin treatment for three consecutive days. After a 10-days hiatus from vancomycin treatments, the animals underwent another round of clindamycin treatment. It’s important to note that there were no further *C. difficile* challenges following the clindamycin treatment in the relapse model. Throughout the study, the animals were closely monitored, and their disease symptoms were assessed using the criteria outlined in the primary infection model.

## Results

### 3.1. The C. difficile JIR8094:: glgC mutant

We generated the *C. difficile* JIR8094::*glgC* strain through the disruption of the first gene, *glgC*, within the glycogen operon. This disruption was achieved using the ClosTron mutagenesis system. It’s worth noting that due to *glgC*’s position as the first gene in the glycogen operon and the methodology of the ClosTron mutagenesis system, all genes within the operon were effectively disrupted. We verified this by employing qRT-PCR to measure the transcript levels of *glgC* and the other downstream genes in the operon, namely *glgD*, *glgA*, *glgP* and *amy*. We prepared RNA from the 8h old JIR8094::*glgC* mutant and the WT and synthesized cDNA to perform qRT-PCR analysis. The Ct values recorded for all five genes in the operon were really high (above 35 cycles) indicating the absence of amplification of transcripts in the JIR8094::*glgC* (Supplemental Figure 2B), while these transcripts could be detected (∼Ct values 20) in the wild-type (WT) strain.

To gain insight into the role of glycogen metabolism under normal growth conditions, we conducted a growth curve comparison between the WT and *C. difficile* JIR8094::*glgC* strains. In TY media, there was no significant difference in growth when assessed by OD600. Additionally, the growth analysis using the Colony Forming Units per milliliter (CFU/mL) count method indicated a similar number of CFUs for *C. difficile* JIR8094::*glgC* in comparison to the parent strain. In the presence of glucose (0.5% w/v), however, the parent strain grew better than the JIR8094::*glgC* mutant.

To confirm that glycogen metabolism was indeed affected in the mutant strain, we measured the intracellular glycogen levels using an amyloglycosidase assay. We supplemented the growth media with either excess glucose or raffinose to induce intracellular glycogen accumulation. The results of the assay showed that raffinose (2%) was a more effective inducer of glycogen accumulation compared to glucose (Figure 3A) where the WT JIR8094 strain rapidly accumulated more glycogen at late exponential phase (8h) and the content started declining at the start of the stationary phase indicating utilization of the glycogen. Glygogen measurement of the JIR8094::*glgC* strain grown in the presence of raffinose detected little to no glycogen in contrast to the WT *C. difficile* (Figure 3B).

**Figure 2.**
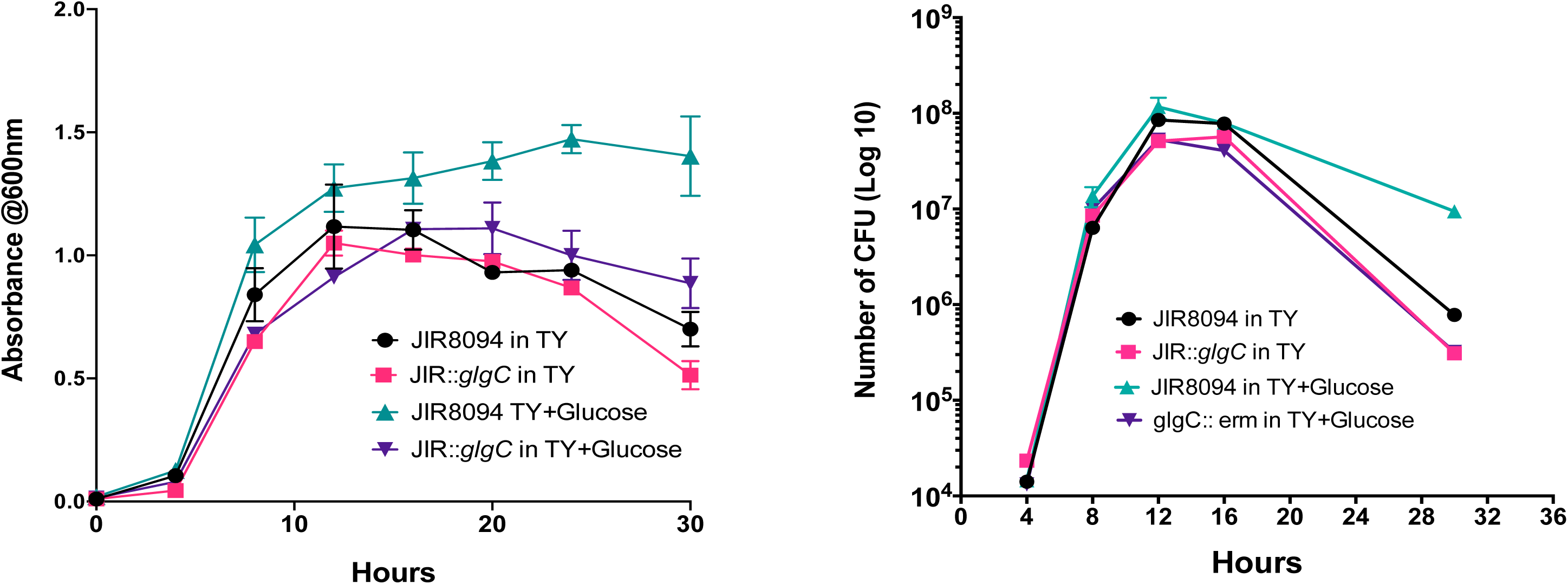
Growth comparison between *C. difficile* JIR8094 and *C. difficile* JIR8094*:: glgC*. (**a**) *C. difficile* parent and mutant strains were grown in TY or TY supplemented with 0.5% glucose. OD_600_ was measured at indicated timepoints. (**b**) Growth measured by CFU/ml count method. Data represents three biological replicates.

**Figure 3.**
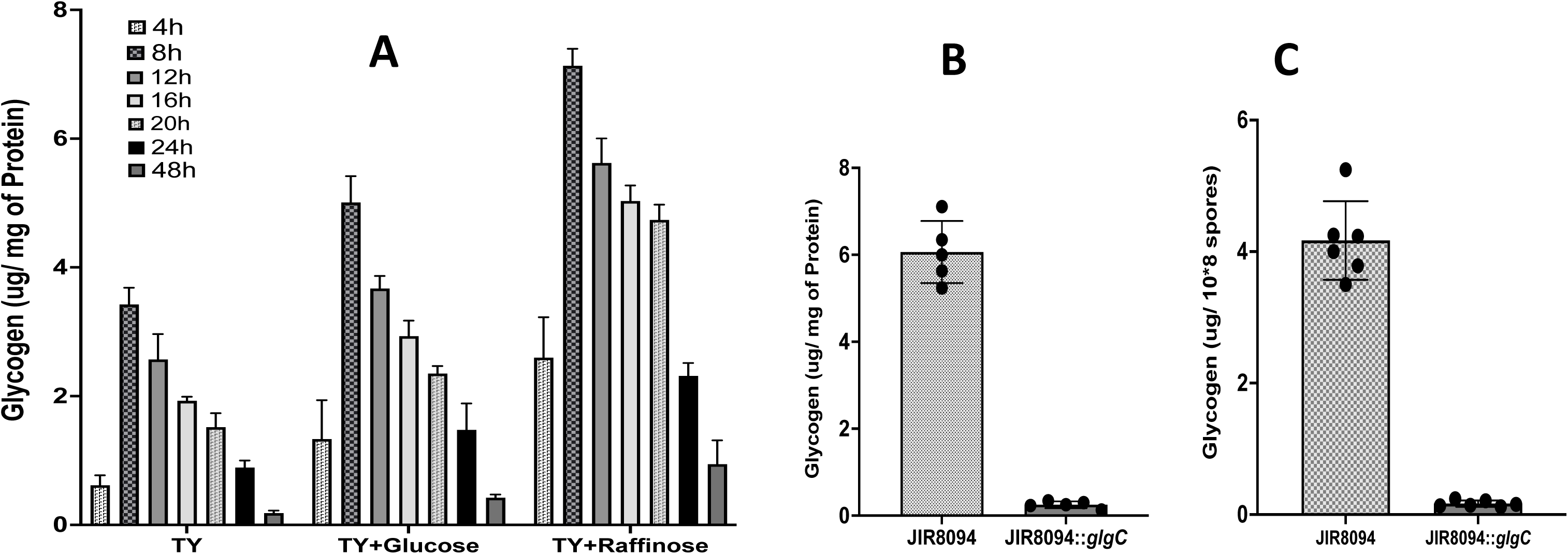
Intracellular glycogen content in the *C. difficile* vegetative cells and spores. **(a)** Intracellular glycogen content of JIR8094 vegetative cells at different timepoints grown in TY or TY supplemented with 0.5% glucose or 2% raffinose. **(b)** Comparison of Intracellular glycogen content between JIR8094 vs JIR8094 *::glgC* vegetative cells at 12h of growth in TY supplemented with 2% Raffinose. **(c)** Comparison of Intracellular glycogen content in spores harvested from JIR8094 and JIR8094 *::glgC* grown in 70:30 media supplemented with 2% raffinose ***=*p*<.001 using Students t-test

### *M*utation in glycogen pathway does not affect the sporulation and toxin production

To understand the effect of glycogen metabolism in *C. difficile* toxin production, we conducted an ELISA-based toxin assay to measure *C. difficile* intracellular toxin levels. The assay result shows that, at a 16-hour time point, *C. difficile* JIR8094:: *glgC* produces slighlty more (Approximately 1.5 times more in mean absorbance level, p<0.01) toxin when compared to the WT strain (Supplemental Figure 3A). Additionally, we compared the sporulation efficiency of the mutant and WT strains using the alcohol sensitivity method. In both these assays, *C. difficile* JIR8094*:: glgC* produced almost similar amount of spores as the parent strain (Supplemental Figure 3B).

While working with spores harvested from the JIR8094::*glgC* mutant strain, we observed a rapid decline in spore viability upon storing at 4°C. Notably, mutant spores lost 100% of their viability within two months of storage at 4°C, whereas spores from the parent strain retained at least 75% of their viability. This observation prompted us to investigate the presence of glycogen in *C. difficile* spores. To do so, we lysed the spores through sonication and measured their glycogen content following the methods outlined in the methods section. Our analysis revealed that *C. difficile* does indeed store glycogen in its spores. Spores harvested from the parent strain contained a significant amount of glycogen, while the glycogen content in the mutant spores was below detectable levels (Figure 3C).

### JIR8094::*glgC* spores are sensitive to chemical and physical disinfecting agents

To assess the significance of glycogen in *C. difficile* spores, we subjected them to various physical and chemical disinfecting agents. When exposed to a 1% bleach solution, spores from the parent strain displayed better resistance, with only about 2% viability loss after 60 minutes, while the JIR8094::*glgC* mutant spores experienced a 90% loss of viability after 60 minutes (Figure 4). Even a short period of exposure to 10% bleach solution proved effective against both strains, but the mutant spores exhibited greater sensitivity. Hydrogen peroxide (3% H_2_O_2_) had a significant impact on spore viability, with the JIR8094::*glgC* mutant spores losing all viability within just 5 minutes of exposure. Notably, sulfuric acid at 1N concentration had no impact on spore viability for either strain, even after 60 minutes of exposure. In contrast, alkali treatment proved lethal to the spores, with the mutant spores showing increased sensitivity compared to those from the parent strain. Except for the acid treatment, the JIR8094::*glgC* mutant spores displayed heightened sensitivity to all other chemical treatments compared to the parent strain, underscoring the essential role of stored glycogen in spore resilience (Figure 4).

**Figure 4.**
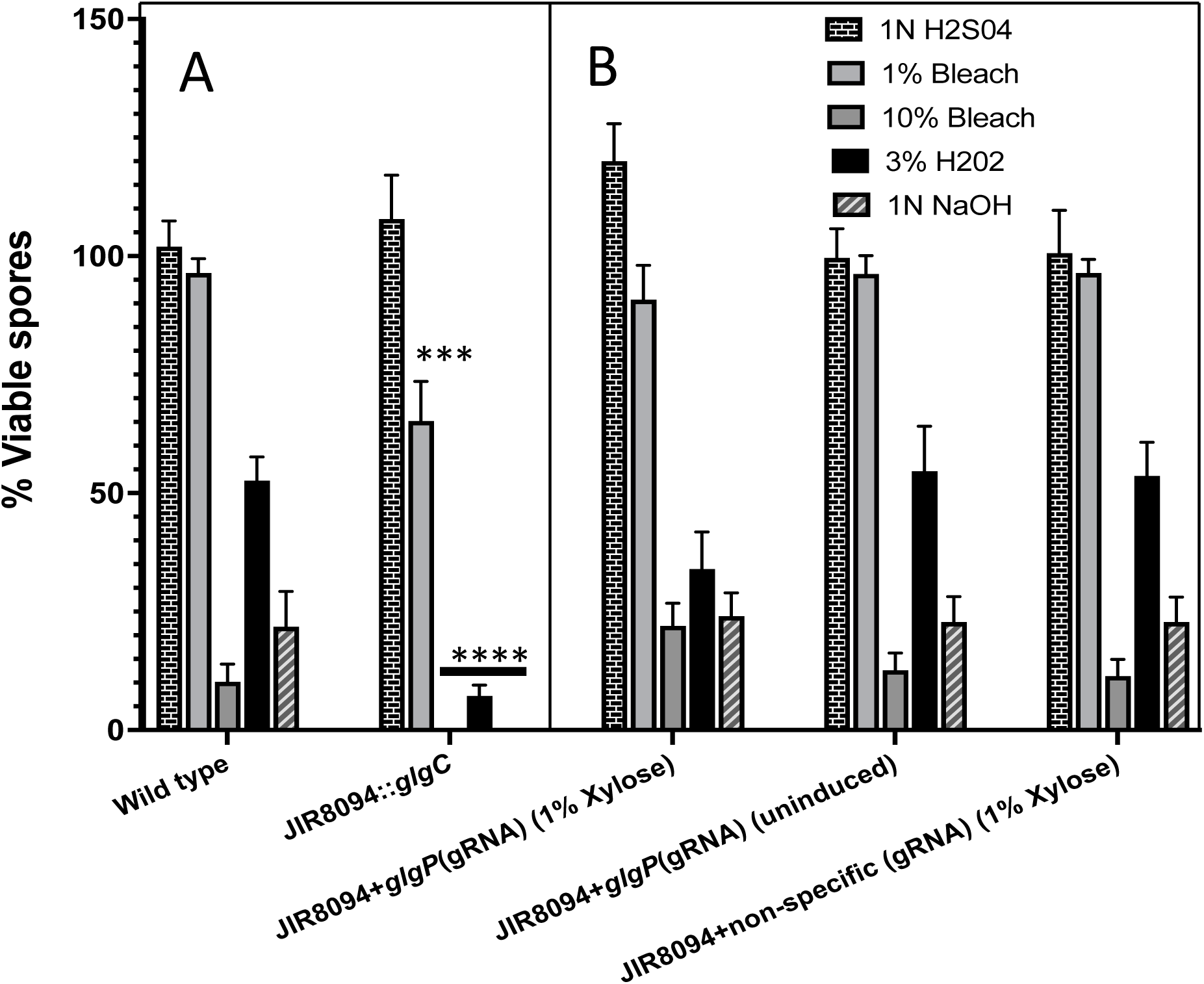
Viability assays of *C. difficile* spores subjected to different stress conditions. Purified spores from *C. difficile* JIR8094, JIR8094*::glgC,* and JIR8094 expressing *glgP*(gRNA) strains along with controls were subjected to 1N H_2_SO_4,_ 1% Bleach, 10% Bleach, 3% H_2_O_2,_ or 1N NaOH treatment. Asterisks (***) indicate a P value of <0.005, and (***) a P value of <0.005 as determined by two-way ANOVA followed by a Dunnett’s multiple comparisons. Comparisons were made with the corresponding treatment of WT strain.

### Only glycogen production, not the utilization is needed for the production of resilient spores

In the *C. difficile* glycogen operon, the first three genes (*glgCDA*) encode enzymes responsible for glycogen synthesis, while the last two genes (*glgP* and *amy*) encode enzymes involved in glycogen breakdown to glucose-1P. To investigate the role of glycogen utilization in spore resilience, we employed a targeted gene knockdown approach using the guide RNA (gRNA) targeting *glgP* gene. This was facilitated through the CRIPRi plasmid pIA33 [19], which carries a gene for an endonuclease-defective Cas9 enzyme under the xylose inducible promoter. This defective Cas9, when guided by the gRNA, binds to the specific DNA region, thereby inhibiting transcription without inducing gene cleavage. This strategy allowed for the selective suppression of glycogen utilization while preserving glycogen synthesis. The plasmid pIA33 containing *glgP*-gRNA (designed using CRISPR-SCAN) was introduced into the JIR8094 strain, with a transconjugant carrying pIA33 with a non-specific gRNA serving as a control.

Initially, we quantified glycogen accumulation in *glgP*-gRNA-induced bacterial cultures, confirming a two-fold increase compared to the uninduced strain and the control strain expressing a non-specific gRNA (Figure 4B). To evaluate the impact on spore resilience, both the test and control strains were cultured on 70:30 medium with thioamphinicol (with and without 1% xylose), incubated for 48 to 72 hours, and the resulting spores were prepared. Our analysis revealed that spores derived from the glgP gRNA-expressing strain exhibited significantly higher resilience to all disinfecting agents compared to the controls (Figure 4B). These findings suggest that accumulated glycogen, rather than its utilization, plays a critical role in spore resilience.

### *C. difficile* spores lacking glycogen has thinner core than the WT spores with stored glycogen

Transmission electron microscopy was performed on spores harvested from the JIR8094 and from the JIR8094::*glgC* mutant strain. One observation was that the core of the spores from the JIR8094::*glgC* mutant strain was smaller with a thicker cortex compared to spores from the parent strain (Figure 5). We measured the radius of the core and the cortex of spores using the imageJ software to calculate the core/cortex ratio (Figure 5). Eight representative images from WT and ten images from the *glgC* mutant were used (supplementary file 1) for this analysis and it was found that the core/cortex ratio is nealry 1.2 fold lesser than the WT spores. Even though the exact location of glycogen is not known at this time, we speculate that the glycogen could have been stored in the core. Otther than providing resilence, it could also be an important nutrient source once the spores germinated into vegetative cells.

**Figure 5.**
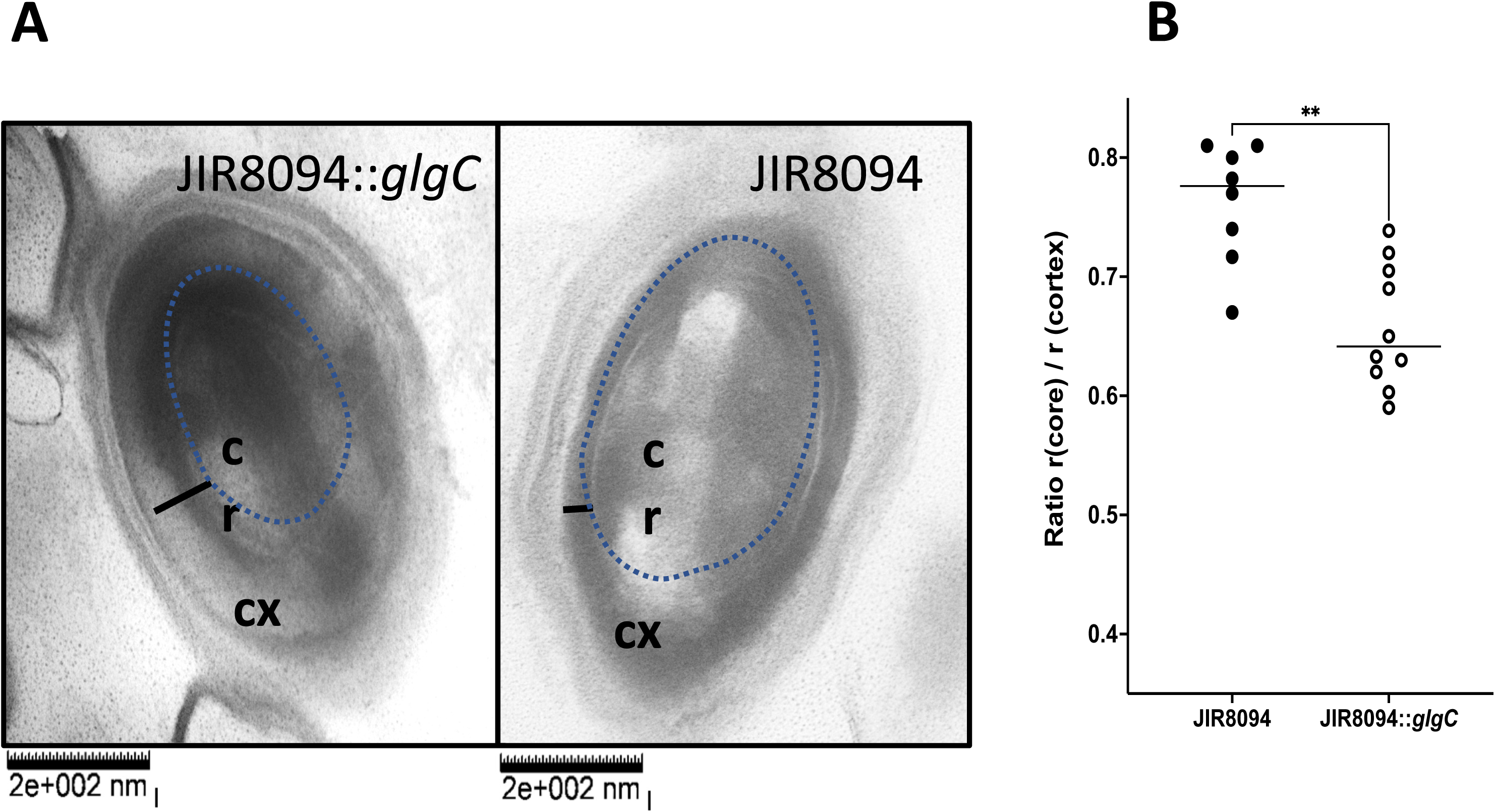
Transmission Electron Microscopic analysis of *C. difficile glgC* mutant spores. (a) TEM image of a spore from the WT (JIR8094) and the JIR8094::*glgC* mutant strain. Cx: Cortex, Cr: core. Dotted lines mark the periphery of the spore cortex. The black bar indicates the thickness of cortex. (b) Comparison of core to cortex ratio in WT and *glgC* mutant (8 spore images from WT and 10 images from JIR8094::*glgC* were used).

### Glycogen is non-essential for initial *C. difficile* colonization and pathogenesis in hamsters, however needed for reinfection

Since the *glgC* mutation was observed to moderately affect *C. difficile* toxin production *in vitro*, we sought to assess the in vivo significance of glycogen production and utilization. Golden Syrian hamsters were orally administered with ∼2000 C. difficile JIR8094 (WT) or *C. difficile* JIR8094::*glgC* and monitored for infections. All hamsters in both the *C. difficile* JIR8094 (WT) infected group and the JIR8094::*glgC* infected group succumbed within 7 days post-infection. Kaplan-Meier survival analysis indicated that the *C. difficile* JIR8094::*glgC* strain exhibited comparable pathogenicity in the animals as the WT strain, suggesting that glycogen production and utilization may not be crucial for C. difficile infection (Figure 6A).

**Figure 6:**
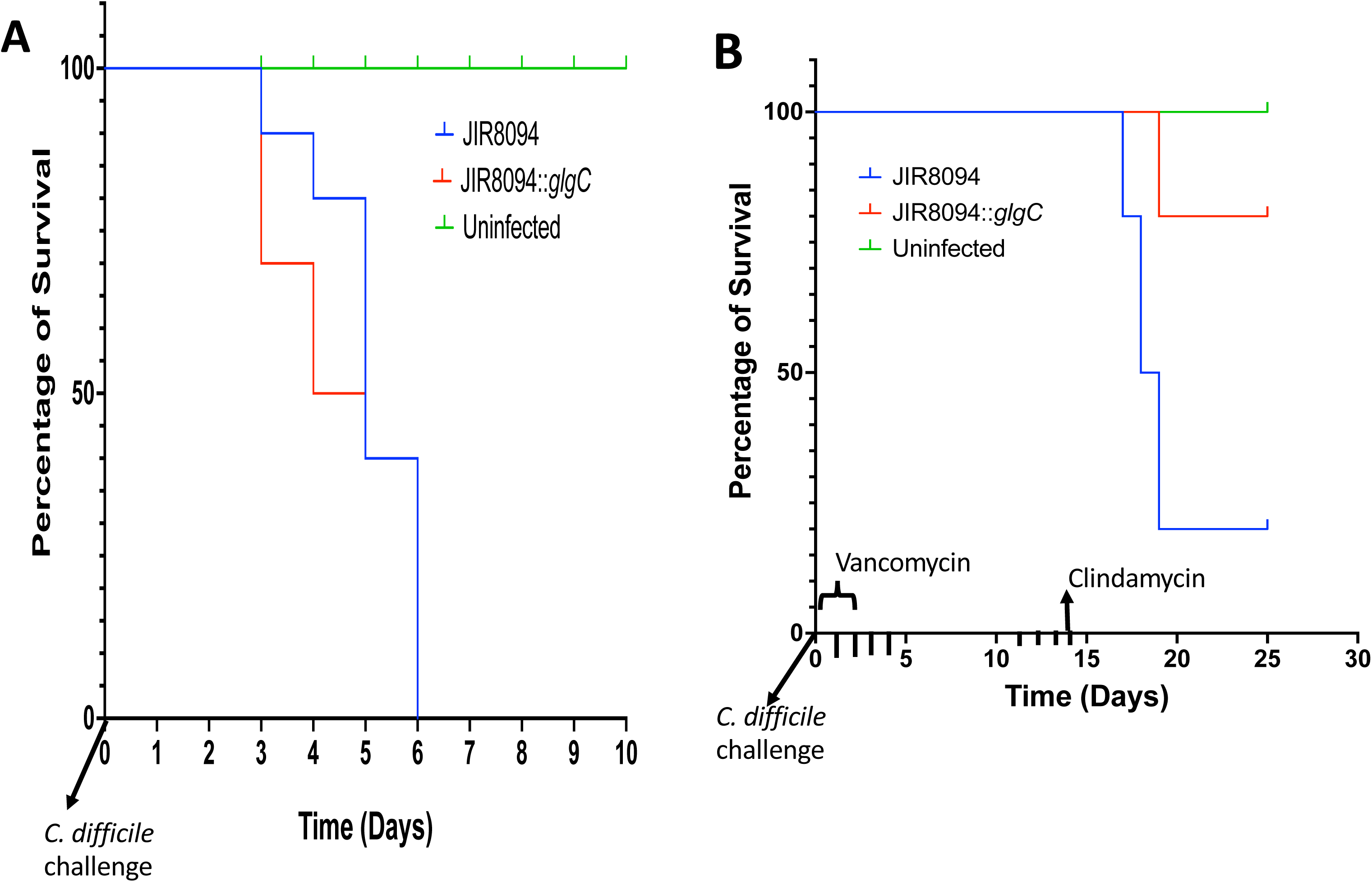
Pathogenicity of JIR8094:: *glgC* in CDI hamster model. Kaplan–Meier survival analysis was conducted to compare the survival rates between the groups, JIR8094 (n=10), JIR8094::*glgC* mutant (n=10), with six uninfected animals serving as controls **(A)** Primary infection model. Clindamycin-treated hamsters were inoculated with either WT or JIR8094:: *glgC* spores. **(B)** In the *C. difficile* relapse model, hamsters were treated with vancomycin for 3 days to clear the primary infection, followed by clindamycin treatment on day 14 to induce a relapse of the infection.

Approximately 50% of *C. difficile* infections result in relapse, and the persistence of infection is linked to sporulation. Since spores isolated from the JIR8094::*glgC* strain demonstrated increased sensitivity to various disinfecting agents, we explored the importance of glycogen storage and utilization in CDI relapse using the hamster relapse infection model. In this model, animals received vancomycin 24 hours post-*C. difficile* challenge for three days. After discontinuation of vancomycin treatments, hamsters received Clindamycin alone and were monitored for CDI symptoms. Results showed that seven out of ten hamsters challenged with JIR8094 (WT) developed the disease, while only one out of ten in the JIR8094::*glgC* challenged group exhibited symptoms (Figure 6B). This suggests that glycogen metabolism is essential for *C. difficile* to survive in the host, potentially as more resilient spores *in vivo*, which could contribute to reinfection.

## Discussion

Glycogen is documented to be a storage carbohydrate for many bacteria [2]. Given how pathogenesis is tightly regulated with nutrition status in *C. difficile*, the ability to store glycogen for use during starvation or other cellular processes can have a significant role in its virulence [4,5,20] [4,5]. Our results indicate that the *glg* operon of *C. difficile* enables glycogen production and significant amount of glycogen is also present in *C. difficile* spores. Our results also show, *C. difficile* JIR8094:: *glgC* produces slightly more toxins compared to WT. However, our *in vivo* study findings indicate very little effect of this increased toxin production in initial *C. difficile* colonization and pathogenicity. A key virulence factor in *C. difficile* is the production of highly resistant exospores. These exospores are the primary method of environmental dissemination as well as recurrent infection occurrence in *C. difficile* patients. Even though *C. difficile* JIR8094:: *glgC* sporulates at similar level as the WT strain, the produced spores are significantly less resilient to exposure to several common stress factors.

In certain spore forming bacteria, glycogen is accumulated at the onset of sporulation and utilized concomitantly during sporulation [21,22]. This would indicate that the energy provided by glycogen degradation is necessary for driving the sporulation process. However, in our case two key observation indicates otherwise. First, the glycogen mutant *C. difficile* was able to produce spores to a similar level that of WT *C. difficile* in the growth conditions tested, and second, significant amount of glycogen was present in the spores produced by WT *C. difficile*. These results indicate that in *C. difficile* sporulation process the primary function of glycogen might not be as energy provider. However, we cannot rule out the role of glycogen in sporulation process in nutrient limited growth conditions. Interestingly, our results of reduced spore viability indicate that glycogen is necessary for proper structural integrity and/or stress tolerance of spores. In this case glycogen itself might be part of a spore structural component, providing resilience. TEM of spores from mutant lacking glycogen showed a smaller core with a thicker cortex compared to the spore with glycogen. The exact reason for this observation is unclear at this point. It is likely that glycogen could be stored in the core to provide immediate energy upon germination. In the Gram-positive spore surface carries several polysaccharides [23]. Glycogen could be used as a precursor for these polysaccharide synthesis pathways and lack of glycogen would result in defective spore surface resulting in reduced stress resistance of glycogen mutant spores. We however ruled out this possibility by demonstrating that glycogen production, but not utilization is needed for the formation of resilient spores.

The role of glycogen in sporulation is best understood in the members of Streptomyces sp. The mycelia forming Gram-positive bacteria. For instance, in *S. antibioticus* and *S. brasiliensis* the onset of sporulation is marked by a significant accumulation of glycogen in sporulating cell [21,22]. However, the accumulated glycogen is degraded during the actual sporulation event in *Streptomyces sp.* [21,22]. Similar to our observation, reducing glycogen content by overexpressing GlgX (a glycogen degrading, debranching enzyme) in *S. venezuelae* resulted in reduced spore viability [24] and also with a distinct color difference between WT spore and *glgX* overexpressed spores indicating a spore structure difference. Glycogen is known to be stored in fungal spores and are part of the spore cytoplasm [25]. The spore specific amylase enzyme (SGA1P) was shown to be important for the glycogen metabolism in the germinating spores in yeast. In the spectroscopy-based germination assay we found the mutant spores lacking glycogen to germinate a little faster than the WT spores, but the difference is not significant (data not shown). This data suggested that the glycogen is not needed for the germination of the spores. However, as we mentioned earlier it could be broken down to glucose-1P and could be utilized for growth once the spores germinated. More work is needed to determine whether glycogen utilizing enzymes are active during the initial stages of growth following spore germination. Besides amino acids, some carbohydrates such as Trehalose and Glycerol are known to be excellent osmotic stress protectors in spores [22]. Glycerol is demonstrated to be rapidly produced from glycogen and whether in *C. difficile* glycogen plays a similar role needs to be investigated.

In the colon, *C. difficile* most probably cycles through nutrient-rich and nutrient-poor conditions. Our *in vivo* study results show that glycogen is not essential for the initial colonization and pathogenesis of *C. difficile* in hamster models which also corroborates our observation of similar growth dynamics between WT and mutant strains in *in vitro* growth condition. So far, our results indicate that glycogen in *C. difficile* is essential for producing viable resilient spores but not for colonization and pathogenesis. In our CDI relapse model, we showed that JIR8094*::glgC* challenged hamsters did not come down with the disease, indicating that spores without glycogen did not survive in the host and didn’t contribute to the relapse of the disease. As dissemination of *C. difficile* occurs primarily through spores, our finding of glycogen being essential for resilient spore production indicate that glycogen playing a significant role in *C. difficile* dissemination. In summary, our study provides a first glimpse of the effects of glycogen metabolism disruption in *C. difficile*.

## Supplementary Figure Legends

**Supplemental Figure 1.**
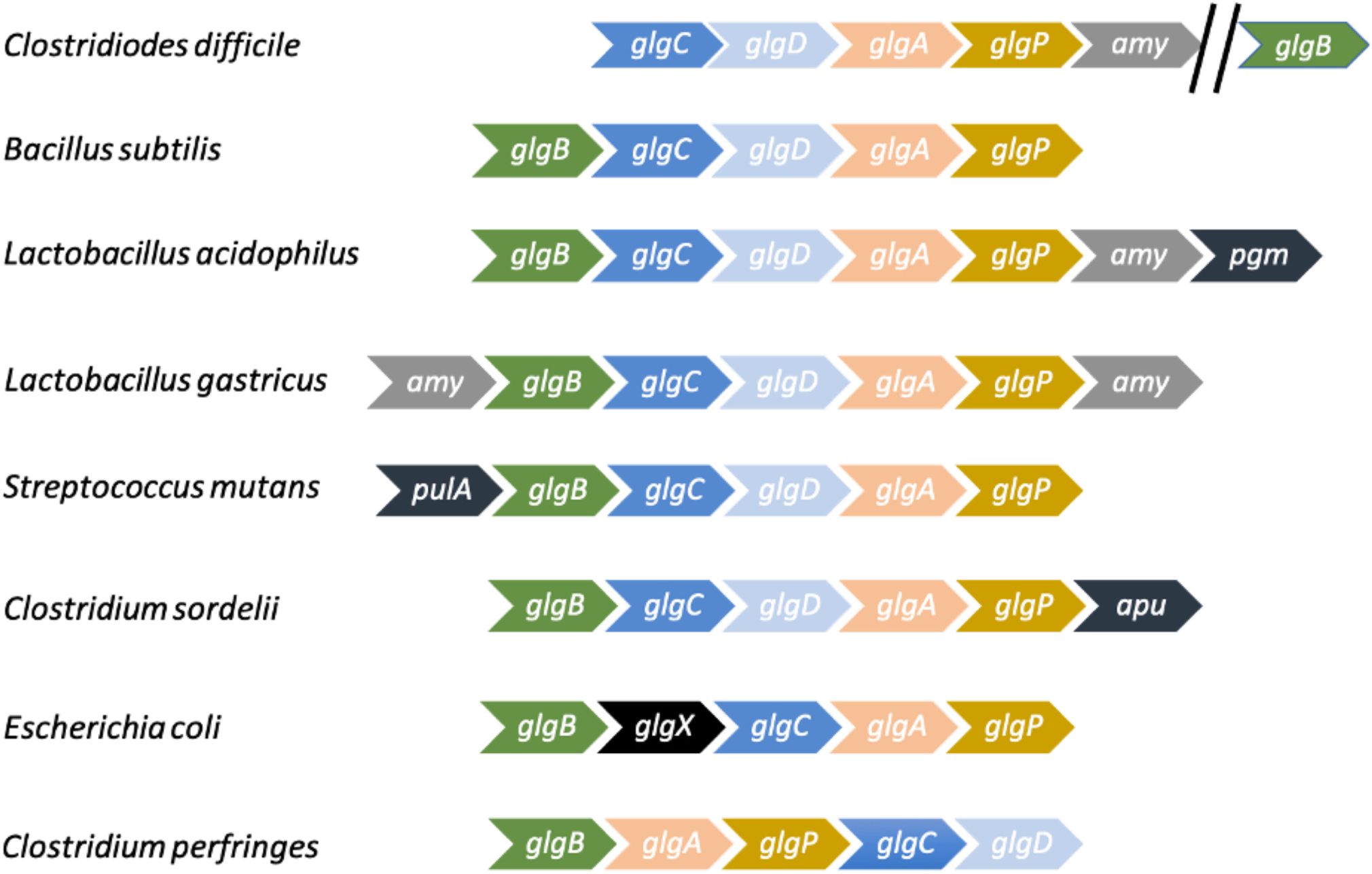
Schematics of glycogen operon in different bacterial species. Orthologous genes are colored same. Glycogen biosynthesis operon is present in many pathogenic and non-pathogenic bacteria. However, the organization of the glycogen biosynthesis and break-down genes varies. Among the examples, *C. difficile* is unique in its location of *glgB* gene (coding for the glycogen branching enzyme) since it in not part of the glycogen operon and is situated at different location in the *C. difficile* genome.

**Supplemental Figure 2.**
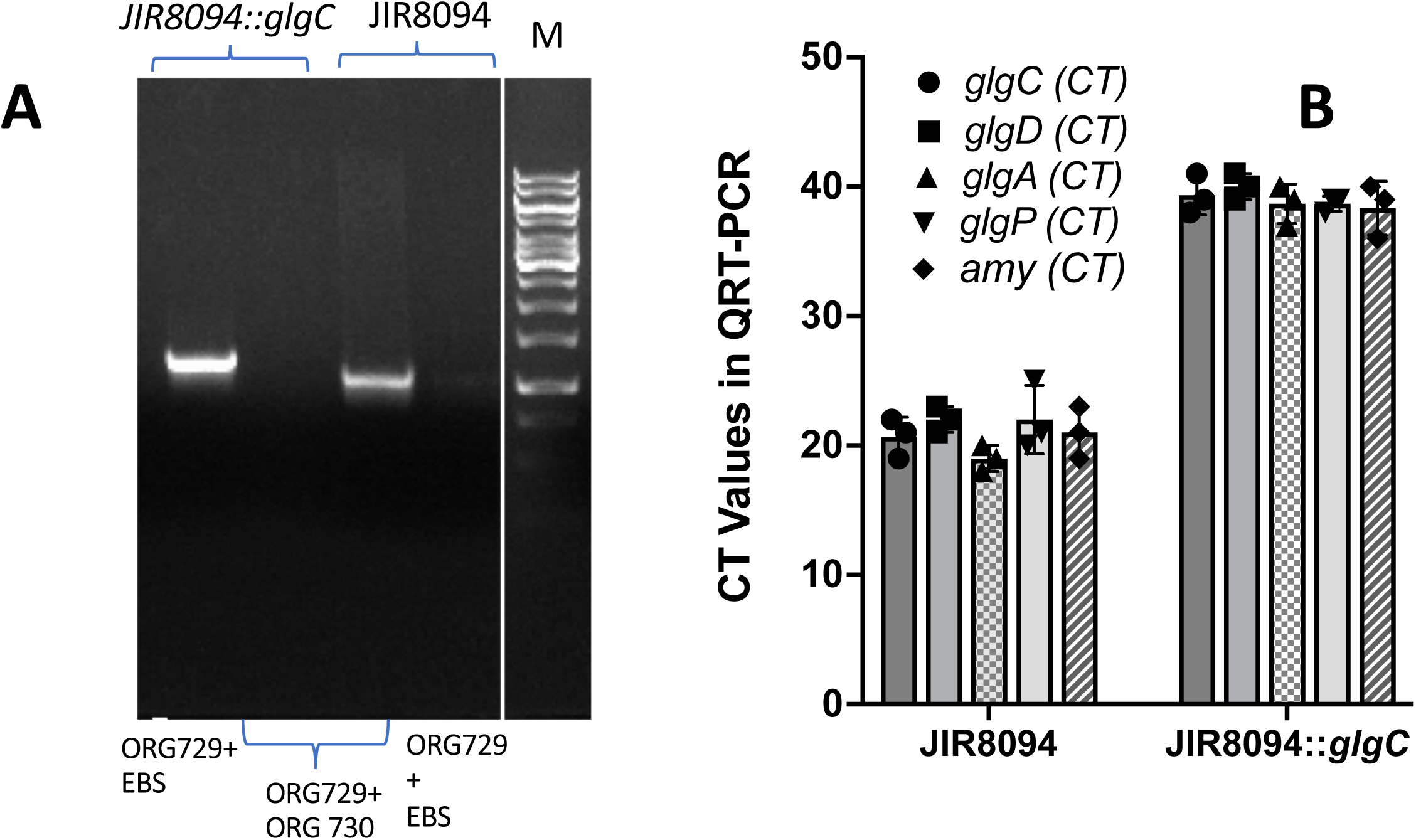
**A.** Agarose gel electrophoresis image of PCR products for verification of JIR8094*::glgC* mutant. An amplification band with intron specific primer EBS universal [EBS(U)], and *glgC* gene specific primer (ORG729) is observed only from the mutant strain but not from the parent strain. An amplification band with gene specific primer pair (ORG729+ORG730) is observed from parent strain but not from the mutant strain since the polymerase is unable to amplify now substantially larger *glgC* gene with the Ll.ltrB intron inserted within. **B.** RT-PCR analysis of *glg* operon genes in WT and *glgC* mutant. All of the *glg* operon transcripts are significantly downregulated in the *glgC* mutant shown by the higher cycle threshold in the *glgC* mutant, which indicates the presence of lower transcript levels.

**Supplemental Figure 3.**
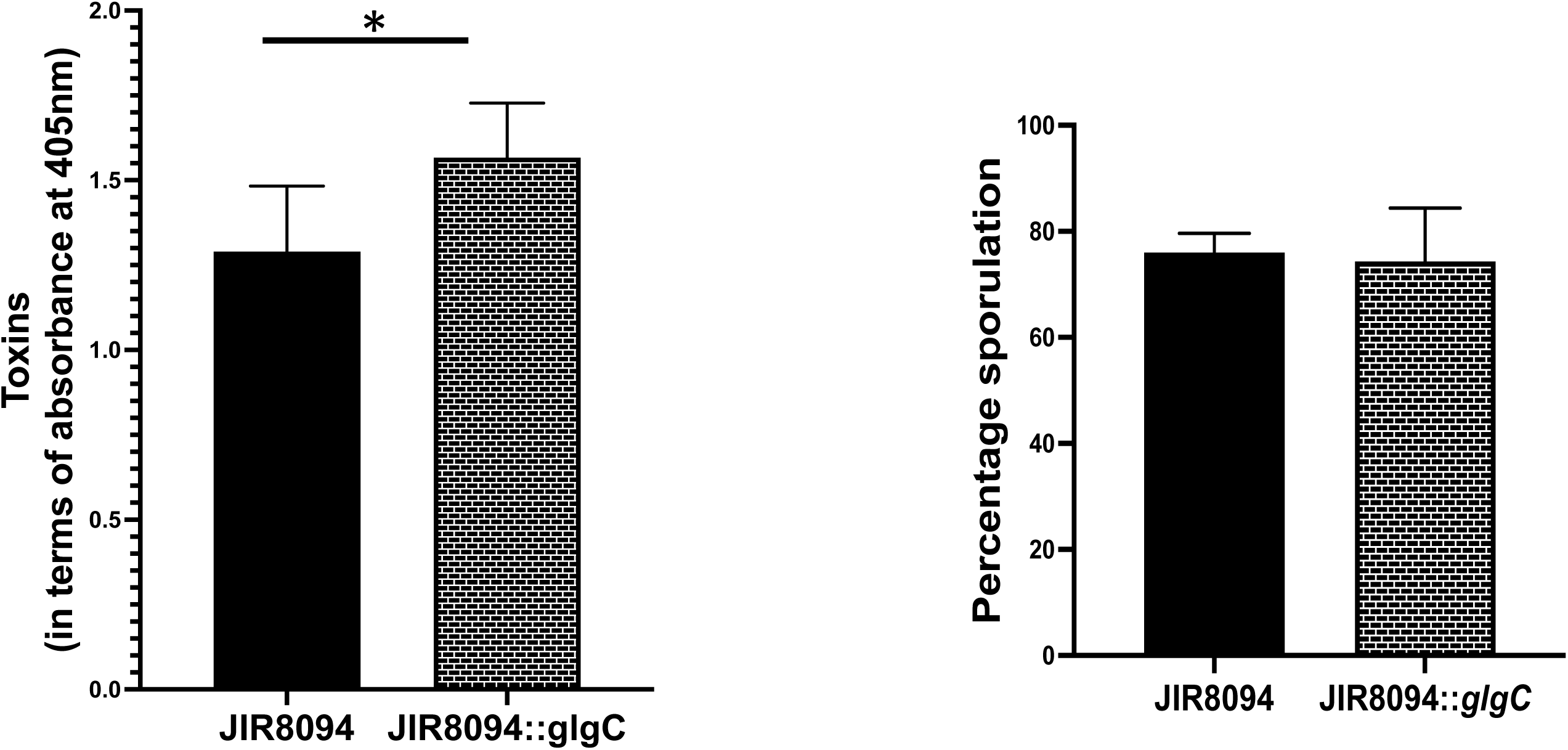
Toxin and sporulation level in the *C. difficile* JIR8094 and *C. difficile* JIR8094*:: glgC:* (**A**) Measurement of cytotoxic Toxin level by ELISA after 16 hours. *P<0.05 using two tailed t-test for means. (**B**) Comparison of Sporulation capacity after 30 Hours between WT and *glgC* mutant strains.

**Supplemental Figure 4:**
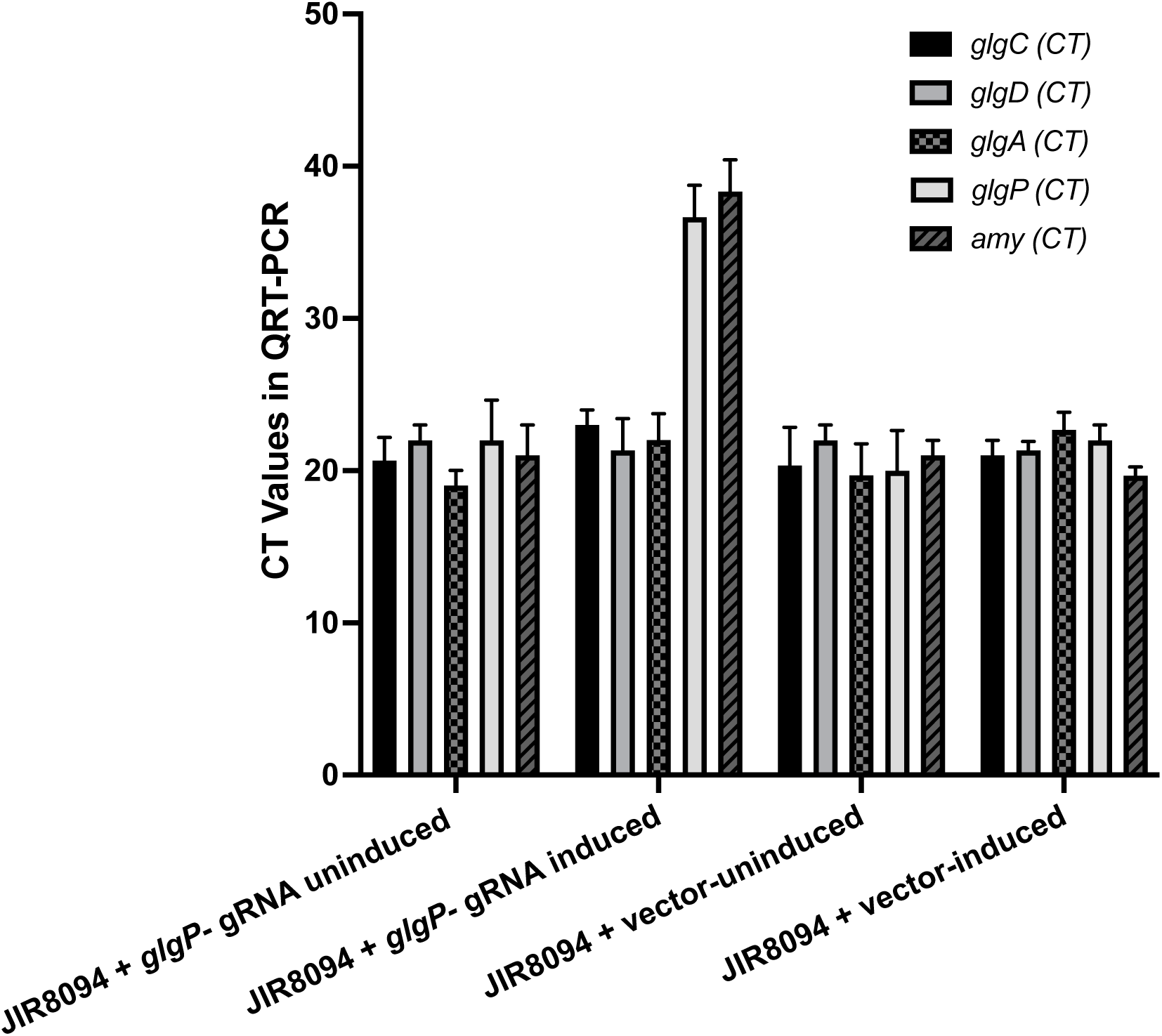
RT-PCR validation of CRISPRi induced inhibition of *glgP* transcription. Increased CT values for *glgP* and *amy* transcripts are observed compared to CRISPi uninduced and vector only control strains. The *amy* gene transcription is downregulated since its situated downstream of *glgP*.

